# ADAR-Sense: an open-access, species-agnostic web tool for automated, user-customisable ADAR-based RNA sensor design

**DOI:** 10.64898/2026.04.16.719097

**Authors:** Henry Sserwadda, Kyemyung Park, Yong-Hee Kim, Hyun Je Kim, Chung-Gyu Park

## Abstract

**Background:** Engineered synthetic RNAs enable cellular control by sensing and responding to intracellular biomolecules. Recently developed sense-edit-switch RNAs (sesRNAs) based on Adenosine Deaminase Acting on RNA (ADAR)—which edits a stop codon to switch on custom payload translation in the presence of a target RNA—are consistently functional across different species. The ability of sesRNAs to couple bespoke payload translation to the presence of cell type-specific transcripts will usher in an era of precise cell-targeted biotechnological interventions.

**Results:** To expedite the generation of sesRNAs, we develop ADAR-Sense—a universal web tool for automated sensor design based on user-defined sensor length, sensor-target RNA mismatch number, mismatch proximity to the ADAR-editable stop codon, and targeted custom element inclusion for improved ADAR recruitment and subsequent payload induction.

**Conclusions:** Compared to current tools, the simplicity and flexibility of ADAR-Sense will streamline the design and screening of sesRNAs in new cell types and conditions, supporting the swift adoption of this sensing platform in both basic and translational research.

## Background

Advances in RNA engineering have facilitated the precise monitoring and programming of cellular states based on intracellular proteins, RNAs, and metabolites, etc^1^. Compared to pre-existing synthetic sensor RNAs, sesRNAs based on adenosine-to-inosine (A-I) editing mediated by adenosine deaminase acting on RNA (ADAR) have demonstrated facile programmability and high dynamic range in various cell types, including mammalian and plant cells^2–4^. SesRNAs contain an in-frame UAG stop codon in a region complementary to a target transcript (target), preventing the ribosome from translating a downstream payload (Fig. 1a). In target-expressing cells, the sesRNA senses the target via base-pairing to form a duplex that recruits ADAR. ADAR edits UAG to UIG, read as UGG by the ribosome, switching on payload translation. The modularity of sesRNAs will enable the reprogramming of specific cell types, thereby accelerating the translation of transcriptomics insights into cutting-edge precision therapeutics.

**Fig. 1:**
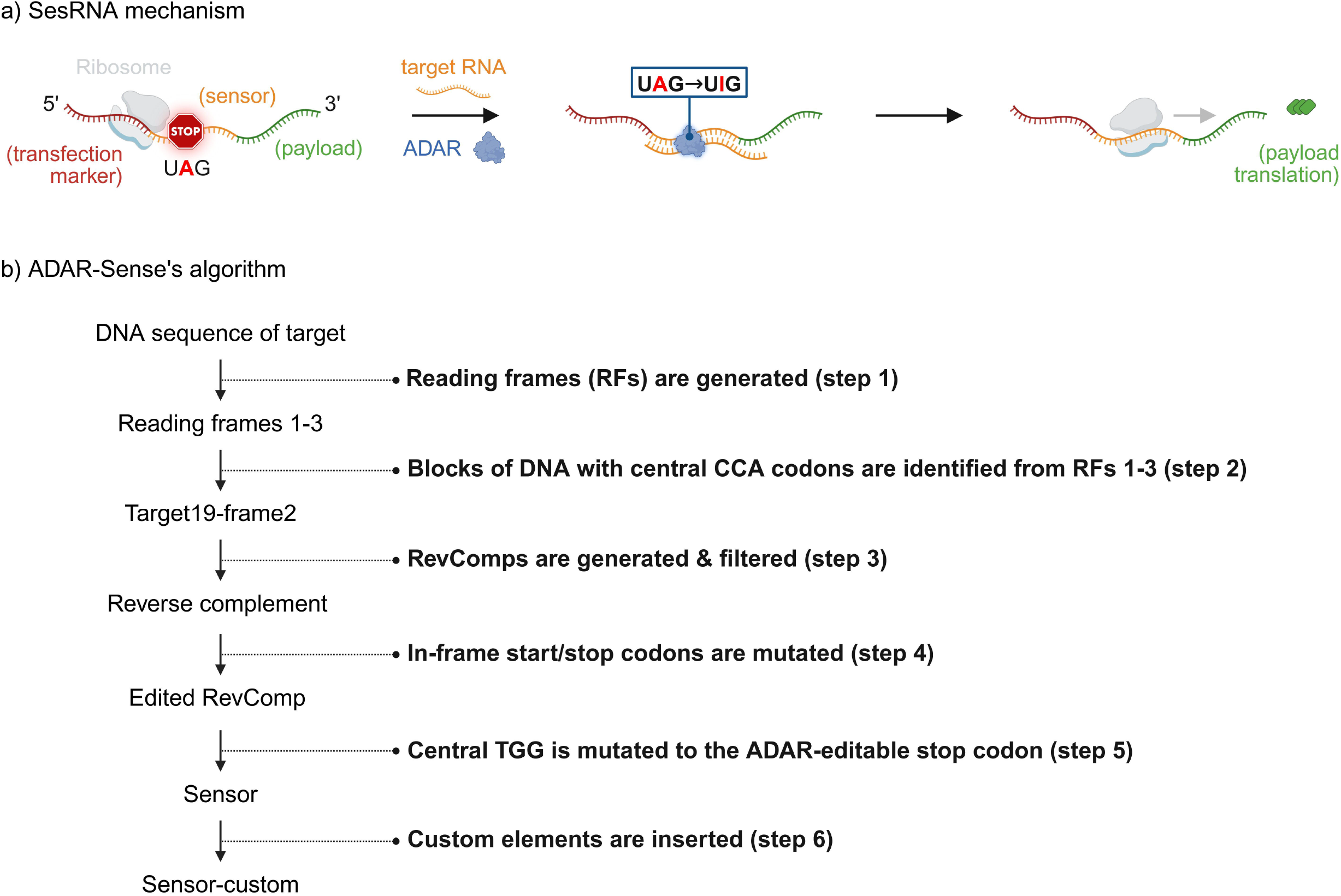
SesRNA mechanism and ADAR-Sense’s features. a) SesRNA encodes a 5’ peptide (which may be a transfection or normalisation marker), a stop codon-embedded sensor region, and a downstream payload. Binding of the target to the sensor results in a duplex which recruits ADAR. ADAR recodes UAG to UIG, enabling payload translation. b) Once the sequence of the target is entered, ADAR-Sense follows six processing steps: reading frame generation, target identification, reverse complement (RevComp) generation and filtering, in-frame start/stop codon mutation, central TGG mutation to TAG, and optional custom element insertion. Fig. 1 was created using BioRender^24^.

Because sesRNAs rely on base-pairing and A-I editing, their performance is influenced by target abundance, target region (i.e., whether target sequences are situated within coding regions, exons, introns, intron-exon junctions, or untranslated regions), ADAR expression level, and the isoform or variant of exogenously supplemented ADAR^2–7^. Other factors such as cell type, sesRNA-target duplex mismatches, sensor region length, sesRNA structural features (e.g., ADAR-recruiting hairpins)^2–8^, and promoter choice for DNA vector-encoded sesRNAs^3^ additionally affect sesRNA performance. To date, cell lines have been used to evaluate the performance of sesRNAs, identifying a range of optimal sensor lengths and structures i.e., 200-350 nucleotides (nt)^2^; 72-153 nt^3^; 252 nt, including four MS2 hairpins^4^; 123 nt, including two MS2 hairpins^6^; 60-81 nt, including one hairpin^7^; and 141 nt^8^. Sun et al. initially screened 66 sesRNAs targeting different regions of hepatitis B virus RNAs, followed by an additional screen of 21 sesRNAs of varying lengths against the identified suitable target region^8^.

Forward, the screening and optimisation of sesRNAs for new experimental contexts, including primary cells of various origins, will require extensive sesRNA design—which is a laborious and error-prone process. Current tools for designing sesRNAs do not accommodate all known species^2,3^, generate reverse complement sequences with in-frame start and stop codons as final outputs^2^, and design sesRNAs of fixed lengths and structures^2–4^ (Table 1). To expedite sesRNA design, we present **A**utomated **D**esign of **A**DAR-based **R**NA **Sens**ors (ADAR-Sense, https://adar-sense.streamlit.app/), a universal web server for generating sesRNAs based on user-defined length, sensor-target mismatch number, mismatch proximity to ADAR-editable UAG, and targeted custom element inclusion.

**Table 1:** ADAR-Sense vs. other sensor design tools.

| Sensor design tool | Automated tool functions and features of designed sensors |  |  |  |  |  |  |
| --- | --- | --- | --- | --- | --- | --- | --- |
|  | Length (flexibility) | Species support | Synthetic target support | Sensor processing | Presence of custom RNA motifs in sensor | add custom RNA motifs | Sensor filtering based on mismatch number & distribution |
| CellREADR<sup>2,9</sup> | ~ 102-300 nucleotides (nt) (fixed range) | Mouse; human | No | Incomplete (contains in-frame start & stop codons) | Absent | No | No |
| RADAR<sup>3,10</sup> | 72 nt, 90 nt (fixed) | Mouse; human; others available upon request | No | Complete | Absent | No | No |
| RADARS<sup>4,11</sup> | 252 nt (fixed) | All | Yes | Complete | Four fixed MS2 loops (at fixed positions) | No | No |
| ADAR-Sense (this work) | ≥ 27 nt (flexible) | All | Yes | Complete | User can add RNA motifs | Yes (at any sensor position) | Yes |

## Implementation

ADAR-Sense is deployed as a standalone Streamit web-based application for automated sensor design. It takes DNA sequences of targets from any source, including natural and synthetic sources, as inputs and generates three reading frames (RFs) in step 1 (Fig. 1b, supplementary file 1). All sequences are processed as DNA for direct use in subsequent molecular cloning experiments. From each RF, DNA chunks of user-defined length with central CCA codons are generated (step 2). Sensor length must be a multiple of three to maintain sesRNA’s frame. The minimum length is set to 27 nt^7^ to enable sensor optimisation in unexplored contexts and to facilitate the design of short sesRNAs suited for scenarios where cargo packaging capacity is limited e.g., with adeno-associated virus-based delivery.

Each target’s reverse complement (RevComp) is then generated, with in-frame start and stop codons colour-coded in green and red, respectively (supplementary file 2). The positions of the in-frame start and stop codons in the generated RevComps are provided using zero-based codon indexing. Because fewer sensor-target mismatches and the absence of mismatches in the vicinity of the ADAR-editable stop codon favour sesRNA function^2^, RevComps are filtered in step 3 to remove those with a user-defined number of in-frame start and stop codons as well as those containing in-frame start and stop codons in the vicinity of the central TGG. The filtering defaults are set to a maximum of five mismatches and a mismatch-free region of 12 nt (4 codons) up- and down-stream of the ADAR-editable stop codon^2,5^ (Fig. 2). To prevent unintended translation initiation and termination, all in-frame start (ATG) and stop (TAG, TAA, TGA) codons are respectively mutated to ATC, TAC, TAC, and TCA^5^. The in-frame start or stop codons may alternatively be substituted as follows: TAG -> TTG, TGG, TCG, TAT, TAC; TGA -> TTA, TCA, TGT, TGG, TGC; TAA -> TTA, TCA, TAT, TAC; and ATG -> TTG, GTG, CTG, ATT, ATA, ATC. To prevent the emergence of mismatched As, which may be subject to unwanted editing by ADAR in the target, caution should be taken to maintain all Ts when introducing mutations in RevComp^5^. The mutation process in step 4 results in undesirable sensor-target mismatches hence the need to pre-emptively perform RevComp filtering in step 3.

**Fig. 2:**
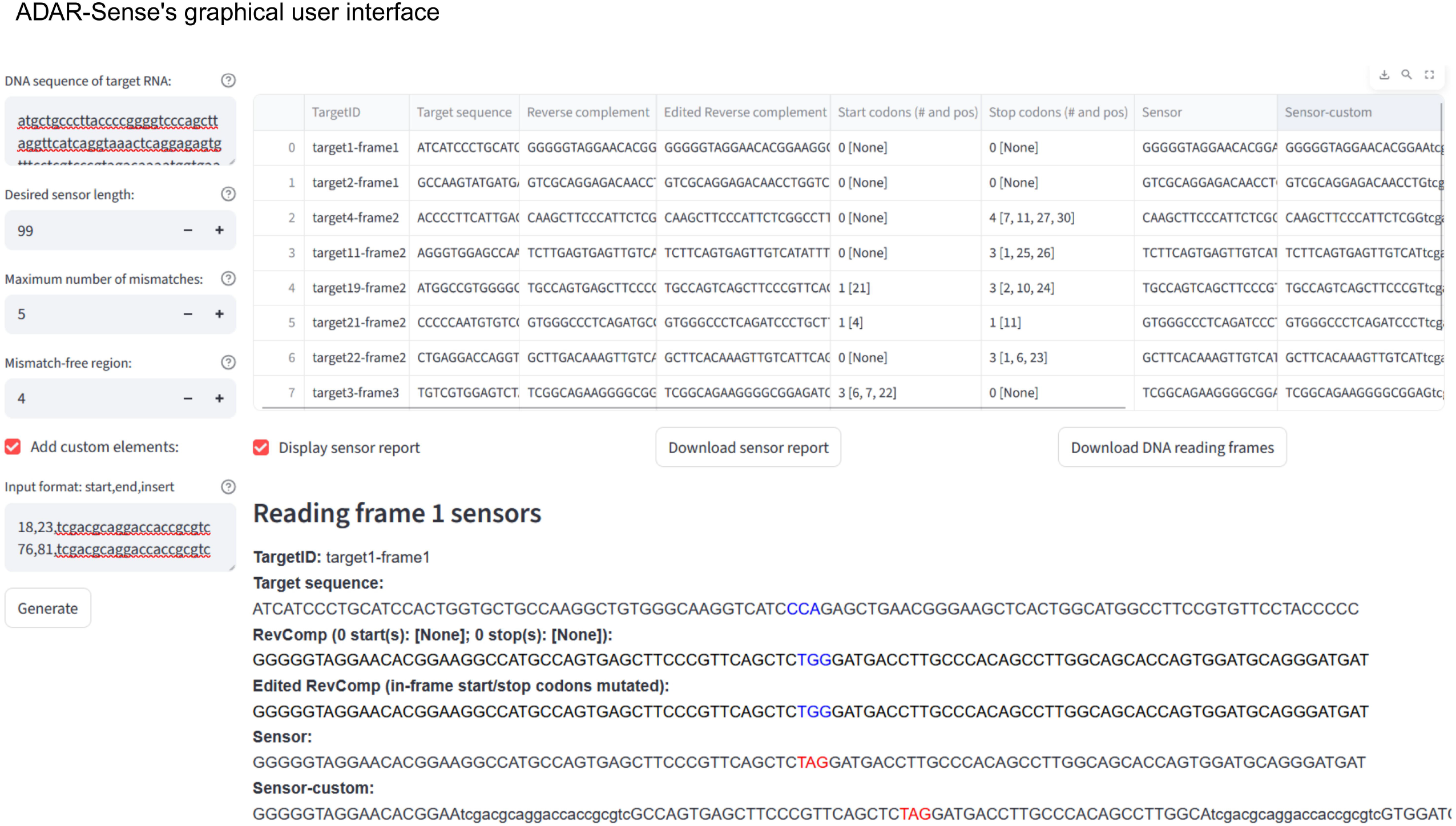
The parameters for mouse Gapdh sensor generation were set to 99 nt, a maximum of five mismatches, and a mismatch-free region of 12 nt (4 codons) up- and down-stream of the ADAR-editable stop codon^5^. The sensor report can be downloaded as an html or csv file (supplementary files 2 and 3). Fig. 2 was created using BioRender^25^.

Finally, the central TGG is mutated to the ADAR-editable stop codon TAG (UAG) to generate ‘sensor’, with the option to introduce new custom elements^3,4,6^ for enhanced sesRNA function. Each custom element and its intended insertion position are entered on a new line; the nt positions are indexed starting from zero (Fig. 2).

A summary of ADAR-Sense’s features in comparison to currently published sensor design tools. The CellREADR portal^2,9^ was inaccessible as of March 2026.

## Results and discussion

To demonstrate the utility of ADAR-Sense, sensors against the coding region of mouse Gapdh^12^ were generated (Fig. 2, Fig. 3a). Sensor length was set to 99 nt, and filtering was performed at ADAR-Sense’s default parameters. Two regions of 6 nt were replaced with two 21-nt-long hairpin motifs^4,13^, maintaining sensor-custom’s ADAR-editable stop codon in frame. Deletions of five to seven nt^4,6^ from the sensor have so far proven sufficient for custom element insertion, as exceedingly short or long deletions are likely to destabilise the sensor-target duplex and compromise sesRNA function. In the target, RevComp, and edited RevComp—all CCA (or TGG), start, and stop codons are respectively highlighted in blue, green, and red in the displayed and downloadable sensor report. Only the ADAR-editable stop codon is coloured red in the final sensor and sensor-custom sequences. The step-by-step colour-coded processing results generated by ADAR-Sense enable users to easily inspect their sensors for errors. The tabulated report contains each sensor’s target region, the (edited) RevComp, the number (#) and positions (pos) of in-frame start and stop codons in the RevComp, and the sensor sequence. See supplementary files 1–3 for the complete sensor processing results.

**Fig. 3:**
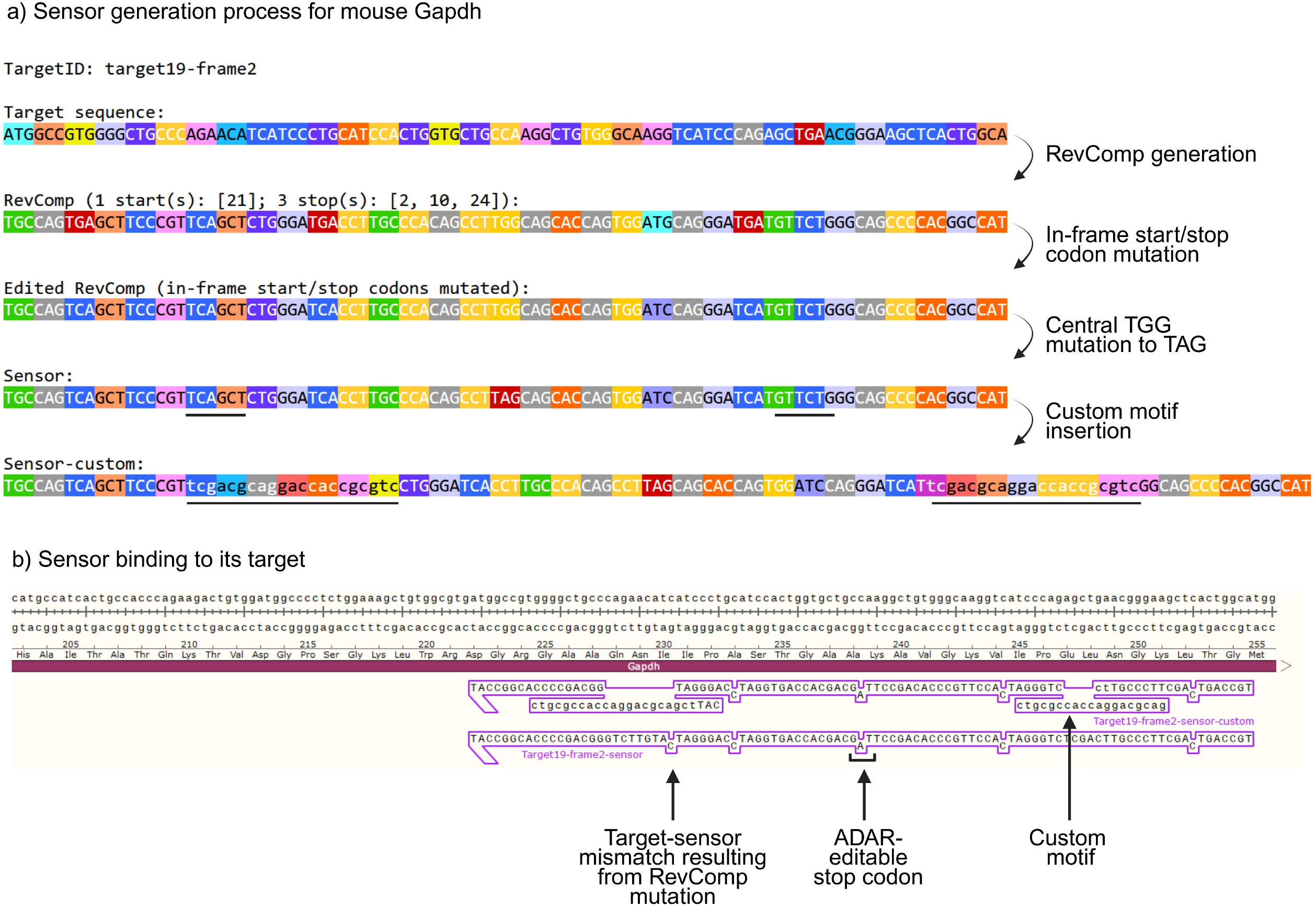
a) Target19-frame2-sensor against mouse Gapdh’s coding region^12^ was designed using the following parameters: 99-nucleotide sensor length, five maximum mismatches, and a mismatch-free region of four codons up- and down-stream of the ADAR-editable stop codon. After target sequence identification, a reverse complement (RevComp) to the target region was generated. RevComp’s in-frame start and stop codons were then eliminated by mutation. The resulting edited RevComp’s central TGG was finally mutated to TAG to generate ‘sensor’. The underlined regions in ‘sensor’ are replaced with ms2_seq5 hairpins^4,13^ to generate ‘sensor-custom’. b) Confirmation of sensor binding to its target using SnapGene Viewer17. Fig. 3 was created using BioRender^26^.

Unlike current design tools^11^, ADAR-Sense enables the user to define the sequence composition, length and insertion position of custom motifs, thus facilitating continued sesRNA optimisation. User-defined elements such as MS2 or BoxB hairpin motifs, naturally edited ADAR substrate motifs, or alu repeat-derived motifs, and synthetic custom motifs, etc^3,4,6,7,14–16^ may be incorporated for improved ADAR recruitment and sensor editing. For sesRNA architectures where the ADAR-editable stop codon is encoded in a hairpin motif at the sensor’s terminal end^7^, the ‘Edited Reverse complement’ sequences (Fig. 2) can be used as the final sensor sequences. For centrally positioned stop codon-encoding hairpin motifs^14^, the custom hairpin motif replaces the region encompassing the ADAR-editable stop codon in the ‘sensor’ sequence to generate ‘sensor-custom’.

During custom element insertion, we recommend making nt deletions and insertions in codons or multiples of three (for example 6 nt may be replaced with a 21-nt motif, as illustrated in Fig. 2). Alternatively, the difference between the deleted and inserted nt must be a multiple of three, for example 23 nt may replace 5 nt (23-5=18, which is a multiple of three). Additionally, custom element insertion must not introduce in-frame start codons or result in multiple in-frame stop codons in each final sensor-custom sequence. This way, frameshifts are avoided, and the translation of the entire sesRNA as a single open RF is maintained. In case of a frame error, ADAR-Sense is programmed to automatically warn the user, hence enabling error-free sensor design. Previous studies have demonstrated that a sensor-target duplex window of at least 41 base pairs (bp), in which the ADAR-editable stop codon is embedded, is required for efficient ADAR catalysis and sesRNA actuation^4,6,7^. Qin PP et al.^7^ showed that flanking the ADAR-editable stop codon with an 11-bp duplex on the 5’ end and a 27- or 39-bp duplex on the 3’ end—achieving respective uninterrupted duplex lengths of 41 (11+UAG+27) and 53 (11+UAG+39) bp—notably improved sesRNA function. Jiang K et al.^4^ and Gayet RV et al.^6^ placed MS2 hairpins outside UAG-embedded duplexes of 51 and 50 bp, respectively, for ADAR recruitment. These findings suggest that maintaining this uninterrupted window during custom element insertion would be good practice for optimal sesRNA design.

The binding of sensors, as reverse primers, to their targets was confirmed using SnapGene Viewer^17^ (Fig. 3b). We noticed that sensors exceeding 250 nt could not be fully entered into SnapGene Viewer and thus recommend binding longer sensors to their targets using more accommodating tools such as Benchling^18^. The sensor-bound target DNA file generated in Benchling can conveniently be exported for visualisation in any preferred DNA analysis software. Users are further advised to confirm that the sensors do not base-pair markedly with other cellular transcripts^5^ using sequence alignment tools such as Basic Local Alignment Search Tool^19,20^. The sesRNAs can finally be cloned into a suitable vector or synthesised in vitro^5^.

## Conclusions

By enabling both novice and expert researchers to generate fully processed sensors based on experiment-specific criteria, ADAR-Sense will fast-track the design, screening, and optimisation of sesRNAs for a wide range of research endeavours. Future in-depth studies on how sensor features, such as secondary structure and sensor-target duplex stability, affect sesRNA performance will enable the screening of functionally superior sesRNAs prior to experimental validation. We anticipate that ADAR-Sense will inspire iterations of sensor design tools that leverage machine learning models trained on vast sesRNA libraries to functionally rank sesRNAs in silico, thereby catalysing innovation across medicine, agriculture, and synthetic biology.

## Availability and requirements

Project name: ADAR-Sense

Project home page: https://adar-sense.streamlit.app/

Operating system(s): Streamlit

Programming language: Python

Other requirements: not applicable

License: End-user license agreement (EULA)^21^

Any restrictions to use by non-academics: ADAR-Sense is available free of charge for academic and non-commercial purposes, with detailed stipulations outlined in the EULA^21^.

## Abbreviations

ADAR: Adenosine Deaminase Acting on RNA
A-I: Adenosine-to-Inosine editing
sesRNA: sense-edit-switch RNA
nt: nucleotide(s)
RF: reading frame
RevComp: reverse complement
bp: base pair(s)

## Ethics approval and consent to participate

Not applicable

## Consent for publication

Not applicable

## Availability of data and materials

The sample DNA sequence of the target used in this study (Fig. 2, Fig. 3) was obtained from the National Center for Biotechnology Information database (accession number NM_001289726.2)^12^. The 21-nt-long ms2_seq5 hairpin sequence (tcgacgcaggaccaccgcgtc) used for custom element insertion (Fig. 2, Fig. 3) was obtained from the ‘find_guides_5avidity.py’ file in the abugoot-lab GitHub repository^4,13^. The rest of the data generated or analysed during this study are included in this published article and its supplementary information files.

## Authors’ contributions

**Henry Sserwadda**: Conceptualisation, Methodology, Software, Writing - original draft, Writing - review and editing, Visualisation. **Kyemyung Park**: Software. **Yong-Hee Kim**: Project administration. **Hyun Je Kim**: Funding acquisition. **Chung-Gyu Park**: Conceptualisation, Resources, Writing - review and editing, Supervision, Project administration, Funding acquisition.

## Acknowledgements

We thank Ji Hwan Moon (Translational Genomics Center, Samsung Medical Center) for valuable feedback on ADAR-Sense. The EULA governing the use of ADAR-Sense^21^ was drafted and revised with legal language refinement assistance from Claude (Anthropic)^22^ and Perplexity (Perplexity AI, Inc)^23^.

## Figure legends

Supplementary file 1: Reading frames generated in Fig. 2.

Supplementary file 2: Sensor report generated in Fig. 2.

Supplementary file 3: Tabulated sensor report generated in Fig. 2.

