## Supplementary files 1-3 for "ADAR-Sense: an open-access, species-agnostic web tool for automated, user-customisable ADAR-based RNA sensor design": Supp1.pdf

### Reading Frames generated from input DNA

**Frame 1:**
ATG CTG CCC TTA CCC CGG GGT CCC AGC TTA GGT TCA TCA GGT AAA CTC AGG AGA GTG TTT CCT CGT CCC GTA GAC AAA ATG GTG AAG GTC GGT GTG AAC GGA TTT GGC CGT ATT GGG CGC CTG GTC ACC AGG GCT GCC ATT TGC AGT GGC AAA GTG GAG ATT GTT GCC ATC AAC GAC CCC TTC
ATT GAC CTC AAC TAC ATG GTC TAC ATG TTC CAG TAT GAC TCC ACT CAC GGC AAA TTC AAC GGC ACA GTC AAG GCC GAG AAT GGG AAG CTT GTC ATC AAC GGG AAG CCC ATC ACC ATC TTC CAG GAG CGA GAC CCC ACT AAC ATC AAA TGG GGT GAG GCC GGT GCT GAG TAT GTC GTG GAG TCT
ACT GGT GTC TTC ACC ACC ATG GAG AAG GCC GGG GCC CAC TTG AAG GGT GGA GCC AAA AGG GTC ATC ATC TCC GCC CCT TCT GCC GAT GCC CCC ATG TTT GTG ATG GGT GTG AAC CAC GAG AAA TAT GAC AAC TCA CTC AAG ATT GTC AGC AAT GCA TCC TGC ACC ACC AAC TGC TTA GCC CCC
CTG GCC AAG GTC ATC CAT GAC AAC TTT GGC ATT GTG GAA GGG CTC ATG ACC ACA GTC CAT GCC ATC ACT GCC ACC CAG AAG ACT GTG GAT GGC CCC TCT GGA AAG CTG TGG CGT GAT GGC CGT GGG GCT GCC CAG AAC ATC ATC CCT GCA TCC ACT GGT GCT GCC AAG GCT GTG GGC AAG GTC
ATC CCA GAG CTG AAC GGG AAG CTC ACT GGC ATG GCC TTC CGT GTT CCT ACC CCC AAT GTG TCC GTC GTG GAT CTG ACG TGC CGC CTG GAG AAA CCT GCC AAG TAT GAT GAC ATC AAG AAG GTG GTG AAG CAG GCA TCT GAG GGC CCA CTG AAG GGC ATC TTG GGC TAC ACT GAG GAC CAG GTT
GTC TCC TGC GAC TTC AAC AGC AAC TCC CAC TCT TCC ACC TTC GAT GCC GGG GCT GGC ATT GCT CTC AAT GAC AAC TTT GTC AAG CTC ATT TCC TGG TAT GAC AAT GAA TAC GGC TAC AGC AAC AGG GTG GTG GAC CTC ATG GCC TAC ATG GCC TCC AAG GAG TAA

**Frame 2:**
TGC TGC CCT TAC CCC GGG GTC CCA GCT TAG GTT CAT CAG GTA AAC TCA GGA GAG TGT TTC CTC GTC CCG TAG ACA AAA TGG TGA AGG TCG GTG TGA ACG GAT TTG GCC GTA TTG GGC GCC TGG TCA CCA GGG CTG CCA TTT GCA GTG GCA AAG TGG AGA TTG TTG CCA TCA ACG ACC CCT TCA
TTG ACC TCA ACT ACA TGG TCT ACA TGT TCC AGT ATG ACT CCA CTC ACG GCA AAT TCA ACG GCA CAG TCA AGG CCG AGA ATG GGA AGC TTG TCA TCA ACG GGA AGC CCA TCA CCA TCT TCC AGG AGC GAG ACC CCA CTA ACA TCA AAT GGG GTG AGG CCG GTG CTG AGT ATG TCG TGG AGT CTA
CTG GTG TCT TCA CCA CCA TGG AGA AGG CCG GGG CCC ACT TGA AGG GTG GAG CCA AAA GGG TCA TCA TCT CCG CCC CTT CTG CCG ATG CCC CCA TGT TTG TGA TGG GTG TGA ACC ACG AGA AAT ATG ACA ACT CAC TCA AGA TTG TCA GCA ATG CAT CCT GCA CCA CCA ACT GCT TAG CCC CCC
TGG CCA AGG TCA TCC ATG ACA ACT TTG GCA TTG TGG AAG GGC TCA TGA CCA CAG TCC ATG CCA TCA CTG CCA CCC AGA AGA CTG TGG ATG GCC CCT CTG GAA AGC TGT GGC GTG ATG GCC GTG GGG CTG CCC AGA ACA TCA TCC CTG CAT CCA CTG GTG CTG CCA AGG CTG TGG GCA AGG TCA
TCC CAG AGC TGA ACG GGA AGC TCA CTG GCA TGG CCT TCC GTG TTC CTA CCC CCA ATG TGT CCG TCG TGG ATC TGA CGT GCC GCC TGG AGA AAC CTG CCA AGT ATG ATG ACA TCA AGA AGG TGG TGA AGC AGG CAT CTG AGG GCC CAC TGA AGG GCA TCT TGG GCT ACA CTG AGG ACC AGG TTG
TCT CCT GCG ACT TCA ACA GCA ACT CCC ACT CTT CCA CCT TCG ATG CCG GGG CTG GCA TTG CTC TCA ATG ACA ACT TTG TCA AGC TCA TTT CCT GGT ATG ACA ATG AAT ACG GCT ACA GCA ACA GGG TGG TGG ACC TCA TGG CCT ACA TGG CCT CCA AGG AGT

**Frame 3:**
GCT GCC CTT ACC CCG GGG TCC CAG CTT AGG TTC ATC AGG TAA ACT CAG GAG AGT GTT TCC TCG TCC CGT AGA CAA AAT GGT GAA GGT CGG TGT GAA CGG ATT TGG CCG TAT TGG GCG CCT GGT CAC CAG GGC TGC CAT TTG CAG TGG CAA AGT GGA GAT TGT TGC CAT CAA CGA CCC CTT CAT
TGA CCT CAA CTA CAT GGT CTA CAT GTT CCA GTA TGA CTC CAC TCA CGG CAA ATT CAA CGG CAC AGT CAA GGC CGA GAA TGG GAA GCT TGT CAT CAA CGG GAA GCC CAT CAC CAT CTT CCA GGA GCG AGA CCC CAC TAA CAT CAA ATG GGG TGA GGC CGG TGC TGA GTA TGT CGT GGA GTC TAC
TGG TGT CTT CAC CAC CAT GGA GAA GGC CGG GGC CCA CTT GAA GGG TGG AGC CAA AAG GGT CAT CAT CTC CGC CCC TTC TGC CGA TGC CCC CAT GTT TGT GAT GGG TGT GAA CCA CGA GAA ATA TGA CAA CTC ACT CAA GAT TGT CAG CAA TGC ATC CTG CAC CAC CAA CTG CTT AGC CCC CCT
GGC CAA GGT CAT CCA TGA CAA CTT TGG CAT TGT GGA AGG GCT CAT GAC CAC AGT CCA TGC CAT CAC TGC CAC CCA GAA GAC TGT GGA TGG CCC CTC TGG AAA GCT GTG GCG TGA TGG CCG TGG GGC TGC CCA GAA CAT CAT CCC TGC ATC CAC TGG TGC TGC CAA GGC TGT GGG CAA GGT CAT
CCC AGA GCT GAA CGG GAA GCT CAC TGG CAT GGC CTT CCG TGT TCC TAC CCC CAA TGT GTC CGT CGT GGA TCT GAC GTG CCG CCT GGA GAA ACC TGC CAA GTA TGA TGA CAT CAA GAA GGT GGT GAA GCA GGC ATC TGA GGG CCC ACT GAA GGG CAT CTT GGG CTA CAC TGA GGA CCA GGT TGT
CTC CTG CGA CTT CAA CAG CAA CTC CCA CTC TTC CAC CTT CGA TGC CGG GGC TGG CAT TGC TCT CAA TGA CAA CTT TGT CAA GCT CAT TTC CTG GTA TGA CAA TGA ATA CGG CTA CAG CAA CAG GGT GGT GGA CCT CAT GGC CTA CAT GGC CTC CAA GGA GTA
