## Supplementary files 1-3 for "ADAR-Sense: an open-access, species-agnostic web tool for automated, user-customisable ADAR-based RNA sensor design": Supp2.pdf

### Reading frame 1 sensors

**TargetID:** target1-frame1
**Target sequence:**
ATCATCCCTGCATCCACTGGTGCTGCCAAGGCTGTGGGCAAGGTCATCCCAGAGCTGAACGGGAAGCTCACTGGCATGGC
**RevComp (0 start(s): [None]; 0 stop(s): [None]):**
GGGGGTAGGAACACGGAAGGCCATGCCAGTGAGCTTCCCGTTCAGCTCTGGGATGACCTTGCCCACAGCCTTGGCAGCA
**Edited RevComp (in-frame start/stop codons mutated):**
GGGGGTAGGAACACGGAAGGCCATGCCAGTGAGCTTCCCGTTCAGCTCTGGGATGACCTTGCCCACAGCCTTGGCAGCA
**Sensor:**
GGGGGTAGGAACACGGAAGGCCATGCCAGTGAGCTTCCCGTTCAGCTCTAGGATGACCTTGCCCACAGCCTTGGCAGCA
**Sensor-custom:**
GGGGGTAGGAACACGGAAtcgacgcaggaccaccgcgtcGCCAGTGAGCTTCCCGTTCAGCTCTAGGATGACCTTGCCCACAGCC

---

**TargetID:** target2-frame1
**Target sequence:**
GCCAAGTATGATGACATCAAGAAGGTGGTGAAGCAGGCATCTGAGGGCCCACTGAAGGGCATCTTGGGCTACACTGAGGA
**RevComp (0 start(s): [None]; 0 stop(s): [None]):**
GTCGCAGGAGACAACCTGGTCCTCAGTGTAGCCCAAGATGCCCTTCAGTGGGCCCTCAGATGCCTGCTTCACCACCTTCT
**Edited RevComp (in-frame start/stop codons mutated):**
GTCGCAGGAGACAACCTGGTCCTCAGTGTAGCCCAAGATGCCCTTCAGTGGGCCCTCAGATGCCTGCTTCACCACCTTCT
**Sensor:**
GTCGCAGGAGACAACCTGGTCCTCAGTGTAGCCCAAGATGCCCTTCAGTAGGCCCTCAGATGCCTGCTTCACCACCTTCT
**Sensor-custom:**
GTCGCAGGAGACAACCTGtcgacgcaggaccaccgcgtcAGTGTAGCCCAAGATGCCCTTCAGTAGGCCCTCAGATGCCTGCTTCA

---

### Reading frame 2 sensors

**TargetID:** target4-frame2
**Target sequence:**
ACCCCTTCATTGACCTCAACTACATGGTCTACATGTTCCAGTATGACTCCACTCACGGCAAATTCAACGGCACAGTCAAGGC
**RevComp (0 start(s): [None]; 4 stop(s): [7, 11, 27, 30]):**
CAAGCTTCCCATTCTCGGCCTTGACTGTGCCGTTGAATTTGCCGTGAGTGGAGTCATACTGGAACATGTAGACCATGTAGTT
**Edited RevComp (in-frame start/stop codons mutated):**
CAAGCTTCCCATTCTCGGCCTTCACTGTGCCGTTCAATTTGCCGTGAGTGGAGTCATACTGGAACATGTAGACCATGTAGTT
**Sensor:**
CAAGCTTCCCATTCTCGGCCTTCACTGTGCCGTTCAATTTGCCGTGAGTAGAGTCATACTGGAACATGTAGACCATGTAGTT
**Sensor-custom:**
CAAGCTTCCCATTCTCGGtcgacgcaggaccaccgcgtcCTGTGCCGTTCAATTTGCCGTGAGTAGAGTCATACTGGAACATGTAGA

---

**TargetID:** target11-frame2
**Target sequence:**
AGGGTGGAGCCAAAAGGGTCATCATCTCCGCCCCTTCTGCCGATGCCCCCATGTTTGTGATGGGTGTGAACCACGAGAAA
**RevComp (0 start(s): [None]; 3 stop(s): [1, 25, 26]):**
TCTTGAGTGAGTTGTCATATTTCTCGTGGTTCACACCCATCACAAACATGGGGGCATCGGCAGAAGGGGCGGAGATGATGA
**Edited RevComp (in-frame start/stop codons mutated):**
TCTTCAGTGAGTTGTCATATTTCTCGTGGTTCACACCCATCACAAACATGGGGGCATCGGCAGAAGGGGCGGAGATCATCA
**Sensor:**
TCTTCAGTGAGTTGTCATATTTCTCGTGGTTCACACCCATCACAAACATAGGGGCATCGGCAGAAGGGGCGGAGATCATCA
**Sensor-custom:**
TCTTCAGTGAGTTGTCATtcgacgcaggaccaccgcgtcCGTGGTTCACACCCATCACAAACATAGGGGCATCGGCAGAAGGGGCG

---

**TargetID:** target19-frame2
**Target sequence:**
ATGGCCGTGGGGCTGCCCAGAACATCATCCCTGCATCCACTGGTGCTGCCAAGGCTGTGGGCAAGGTCATCCCAGAGCT
**RevComp (1 start(s): [21]; 3 stop(s): [2, 10, 24]):**
TGCCAGTGAGCTTCCCGTTCAGCTCTGGGATGACCTTGCCCACAGCCTTGGCAGCACCAGTGGATGCAGGGATGATGTTC
**Edited RevComp (in-frame start/stop codons mutated):**
TGCCAGTCAGCTTCCCGTTCAGCTCTGGGATCACCTTGCCCACAGCCTTGGCAGCACCAGTGGATCCAGGGATCATGTTC
**Sensor:**
TGCCAGTCAGCTTCCCGTTCAGCTCTGGGATCACCTTGCCCACAGCCTAGCAGCACCAGTGGATCCAGGGATCATGTTCT
**Sensor-custom:**
TGCCAGTCAGCTTCCCGTtcgacgcaggaccaccgcgtcCTGGGATCACCTTGCCCACAGCCTTAGCAGCACCAGTGGATCCAGGG

---

**TargetID:** target21-frame2
**Target sequence:**
CCCCCAATGTGTCCGTCGTGGATCTGACGTGCCGCCTGGAGAAACCTGCCAAGTATGATGACATCAAGAAGGTGGTGAAG
**RevComp (1 start(s): [4]; 1 stop(s): [11]):**
GTGGGCCCTCAGATGCCTGCTTCACCACCTTCTTGATGTCATCATACTTGGCAGGTTTCTCCAGGCGGCACGTCAGATCCA
**Edited RevComp (in-frame start/stop codons mutated):**
GTGGGCCCTCAGATCCCTGCTTCACCACCTTCTTCATGTCATCATACTTGGCAGGTTTCTCCAGGCGGCACGTCAGATCCA
**Sensor:**
GTGGGCCCTCAGATCCCTGCTTCACCACCTTCTTCATGTCATCATACTTAGCAGGTTTCTCCAGGCGGCACGTCAGATCCA
**Sensor-custom:**
GTGGGCCCTCAGATCCCTtcgacgcaggaccaccgcgtcCCACCTTCTTCATGTCATCATACTTAGCAGGTTTCTCCAGGCGGCACG

---

**TargetID:** target22-frame2

**Target sequence:**

CTGAGGACCAGGTTGTCTCCTGCGACTTCAACAGCAACTCCCACTCTTCCACCTTCGATGCCGGGGCTGGCATTGCTCTC

**RevComp (0 start(s): [None]; 3 stop(s): [1, 6, 23]):**

GCTTGACAAAGTTGTCATTGAGAGCAATGCCAGCCCCGGCATCGAAGGTGGAAGAGTGGGAGTTGCTGTTGAAGTCGCAG

**Edited RevComp (in-frame start/stop codons mutated):**

GCTTCACAAAGTTGTCATTCAGAGCAATGCCAGCCCCGGCATCGAAGGTGGAAGAGTGGGAGTTGCTGTTCAAGTCGCAG

**Sensor:**

GCTTCACAAAGTTGTCATTCAGAGCAATGCCAGCCCCGGCATCGAAGGTAGAAGAGTGGGAGTTGCTGTTCAAGTCGCAG

**Sensor-custom:**

GCTTCACAAAGTTGTCATtcgacgcaggaccaccgcgtcCAATGCCAGCCCCGGCATCGAAGGTAGAAGAGTGGGAGTTGCTGTTC

---

### Reading frame 3 sensors

**TargetID:** target3-frame3

**Target sequence:**

TGTCGTGGAGTCTACTGGTGTCTTCACCACCATGGAGAAGGCCGGGGCCCACTTGAAGGGTGGAGCCAAAAGGGTCATC

**RevComp (3 start(s): [6, 7, 22]; 0 stop(s): [None]):**

TCGGCAGAAGGGGCGGAGATGATGACCCTTTTGGCTCCACCCTTCAAGTGGGCCCCGGCCTTCTCCATGGTGGTGAAGA

**Edited RevComp (in-frame start/stop codons mutated):**

TCGGCAGAAGGGGCGGAGATCATCACCCTTTTGGCTCCACCCTTCAAGTGGGCCCCGGCCTTCTCCATCGTGGTGAAGAC

**Sensor:**

TCGGCAGAAGGGGCGGAGATCATCACCCTTTTGGCTCCACCCTTCAAGTAGGCCCCGGCCTTCTCCATCGTGGTGAAGAC

**Sensor-custom:**

TCGGCAGAAGGGGCGGAGtcgacgcaggaccaccgcgtcACCCTTTTGGCTCCACCCTTCAAGTAGGCCCCGGCCTTCTCCATCG

---

**TargetID:** target4-frame3

**Target sequence:**

CATCTCCGCCCCTTCTGCCGATGCCCCCATGTTTGTGATGGGTGTGAACCACGAGAAATATGACAACTCACTCAAGATTGT

**RevComp (2 start(s): [23, 32]; 0 stop(s): [None]):**

GTGCAGGATGCATTGCTGACAATCTTGAGTGAGTTGTCATATTTCTCGTGGTTCACACCCATCACAAACATGGGGGCATCG

**Edited RevComp (in-frame start/stop codons mutated):**

GTGCAGGATGCATTGCTGACAATCTTGAGTGAGTTGTCATATTTCTCGTGGTTCACACCCATCACAAACATCGGGGCATCGG

**Sensor:**

GTGCAGGATGCATTGCTGACAATCTTGAGTGAGTTGTCATATTTCTCGTAGTTCACACCCATCACAAACATCGGGGCATCGG

**Sensor-custom:**

GTGCAGGATGCATTGCTGtcgacgcaggaccaccgcgtcTTGAGTGAGTTGTCATATTTCTCGTAGTTCACACCCATCACAAACATCG

---

**TargetID:** target10-frame3

**Target sequence:**

GGGCTACACTGAGGACCAGGTTGTCTCCTGCGACTTCAACAGCAACTCCCACTCTTCCACCTTCGATGCCGGGGCTGGCA

**RevComp (1 start(s): [6]; 1 stop(s): [31]):**

AAGTTGTCATTGAGAGCAATGCCAGCCCCGGCATCGAAGGTGGAAGAGTGGGAGTTGCTGTTGAAGTCGCAGGAGACAA

**Edited RevComp (in-frame start/stop codons mutated):**

AAGTTGTCATTGAGAGCAATCCCAGCCCCGGCATCGAAGGTGGAAGAGTGGGAGTTGCTGTTGAAGTCGCAGGAGACAA

**Sensor:**

AAGTTGTCATTGAGAGCAATCCCAGCCCCGGCATCGAAGGTGGAAGAGTAGGAGTTGCTGTTGAAGTCGCAGGAGACAAC

**Sensor-custom:**

AAGTTGTCATTGAGAGCAtcgacgcaggaccaccgcgtcGCCCCGGCATCGAAGGTGGAAGAGTAGGAGTTGCTGTTGAAGTCGCA
